# Attention-wise masked graph contrastive learning for predicting molecular property

**DOI:** 10.1101/2022.05.08.491075

**Authors:** Hui Liu, Yibiao Huang, Xuejun Liu, Lei Deng

## Abstract

**Motivation:** Accurate and efficient prediction of the molecular property is one of the fundamental problems in drug research and development. Recent advancements in representation learning have been shown to greatly improve the performance of molecular property prediction. However, due to limited labeled data, supervised learning-based molecular representation algorithms can only search limited chemical space and suffer from poor generalizability.

**Results:** In this work, we proposed a self-supervised learning method, ATMOL, for molecular representation learning and properties prediction. We developed a novel molecular graph augmentation strategy, referred to as attention-wise graph masking, to generate challenging positive samples for contrastive learning. We adopted the graph attention network (GAT) as the molecular graph encoder, and leveraged the learned attention weights as masking guidance to generate molecular augmentation graphs. By minimization of the contrastive loss between original graph and augmented graph, our model can capture important molecular structure and higher-order semantic information. Extensive experiments showed that our attention-wise graph mask contrastive learning exhibited state-of-the-art performance in a couple of downstream molecular property prediction tasks. We also verified that our model pretrained on larger scale of unlabeled data improved the generalization of learned molecular representation. Moreover, visualization of the attention heatmaps showed meaningful patterns indicative of atoms and atomic groups important to specific molecular property.

## Introduction

The physicochemical properties of molecules, such as water solubility, lipophilicity, membrane permeability and degree of dissociation, are of great importance to the screening of leading compounds in drug development. As traditional wet-lab experiments is time-consuming and labor-intensive, it is impossible to cover hundred millions of candidate molecules (1). So, many *in silico* methods have been proposed to predict molecular properties, and these methods greatly promoted the efficiency of drug development and return on investment (2).

Molecular representation is crucial to identify various physicochemical properties of molecules (3–5). The feature engineering-based chemical fingerprint, in which each bit represents the absence or presence of a certain biochemical property or substructure, transforms the structural information or properties to a fixed-length vector (6). For example, PubChem fingerprint (7) and extended connectivity fingerprints (ECFPs) (8) are frequently used molecular representations. However, most chemical fingerprints rely on domain knowledge and contain only task-specific information, which often lead to limited performance when applied to down-stream tasks. In recent years, deep learning has achieved remarkable success in natural language processing (NLP) (9), computer vision (CV) (10), and graph structure prediction (11). Many studies have applied deep learning to chemical modeling (12–17), and drug discovery (10, 18, 19), etc. However, the performance of fully-supervised deep learning depends on a large amount of manually labeled samples (13, 20), such as molecules with known properties in our case. When applied to small-size dataset, the fully-supervised model is vulnerable to overfitting and poor generalizability.

In recent years, self-supervised learning has caught much attention because of its better generalizability achieved in multiple fields. Self-supervised learning first run pretraining on large-scale unlabeled dataset to derive latent representation (embedding) (9, 10, 21), and then transfer to downstream tasks to obtain better robustness (22, 23). For molecular property prediction task, a few self-supervised methods have been proposed to learn molecular representation (24–29). These methods fall roughly into two categories: generation-based methods and contrastive learning-based methods. The generative methods learned embedding by establishing specific pretext tasks that encourage the encoder to extract high-order structural information. For example, MG-BERT learned to predict the masked atomic (24), by integrating the local message passing mechanism of graph neural networks (GNNs) into the powerful BERT model (25) to enhance representation learning from molecular graphs. MolGPT (26) trained a transformer-decoder model for the next token prediction task using masked self-attention to generate novel molecules. Contrastive learning encourages augmentations (contrastive views) of the same molecules to have more similar embeddings compared to those generated from different molecules. For example, MolCLR (27) proposed three different graph augmentation by masking nodes or edges or subgraphs, and then maximize the agreement of the augmentations from same molecule while minimizing the agreement of different molecules. CSGNN (28) designed a deep mix-hop graph neural network to capture higher-order dependencies and introduced a self-supervised contrastive learning framework.

MolGNet (29) used both paired subgraph recognition (PSD) and attribute masking (AttrMasking) to achieve node-level and graph-level pre-training, which was shown to improve the ability to extract feature from molecular graphs.

The performance of contrastive representation learning often relies on the quality of augmented views. Current methods generated contrastive views by randomly masking some nodes and edges [28]. However, the way of random masking cannot guide the encoder to detect most important substructure. In this work, we proposed an attention-wise contrastive learning framework for molecular representation and property prediction. Specifically, we first constructed the molecular graph from SMILES, and used the graph attention network (GAT) as encoder to transform molecular graph into latent representation. Next, we leveraged the attention weights of nodes and edges learned by GAT to generate augmentation graph, by masking a percentage of nodes or edges according to their attention weights. By minimization of the contrastive loss between original graph and augmented graph, our model was driven to capture important substructure and thus produce concise and informative molecular representation. Our extensive experiments showed that the molecular representations learned by our method exhibited state-of-the-art performance in various downstream molecular property prediction tasks. Performance comparison verified our method outperformed competitive methods. Moreover, we explored the interpretability and found the attention wights revealed importance patterns of important substructure.

## Materials and methods

### Data source

We downloaded two molecule sets from ZINC database (substance channel). One is the *in vitro* set, which includes 306,347 unique substances reported or inferred to be bioactive at 10 *μM* or better in direct binding assays. All molecules in the *in vitro* set were used in our evaluation experiments. The other set was built from ZINC’s *now* set, which include all in-stock and agent substances for immediate delivery. This dataset includes 9,814,569 unique molecules. For molecule, the open-source tool RDkit (30) was used to transform its SMILES descriptor to molecular graph, in which a node represents an atom and edge represent a chemical bond. The molecular graphs were ready as input of graph attention network.

For performance evaluation on downstream tasks, we chose 7 datasets from MoleculeNet (31), which has established more than forty molecular property prediction tasks. Table 1 showed the total number of molecules in each dataset, as well as the number of subtasks. The 7 datasets covered different molecular properties, including membrane permeability, toxicity and bioactivity. For each dataset, we used the scaffold split from DeepChem (32) to create an 80/10/10 train/valid/test subset. Rather than random split, scaffold-split divided the molecules based on their substructures, making the prediction task more challenging yet realistic.

**Table 1.** Seven datasets representing prediction tasks of different molecular properties

| DataSet | Molecule number | Task number |
| --- | --- | --- |
| BBBP | 2,039 | 1 |
| BACE | 1,513 | 1 |
| HIV | 41,127 | 1 |
| ClinTox | 1,478 | 2 |
| Tox21 | 7,831 | 12 |
| SIDER | 1,478 | 27 |
| MUV | 93,087 | 17 |

## Method

### ATMOL framework

Our framework consisted of two stages: pre-training and transfer learning. As shown in Figure 1, we first performed contrastive learning on large-scale unlabeled datasets to obtain molecular representations, and then applied transfer learning to predict molecular properties. The molecular graph was taken as input, and mapped to latent space by GAT encoder. Meanwhile, an attention-wise masking module closely tracked the GAT encoder is developed to use the attention scores to produce augmented view (masked graph) by masking some nodes or edges. We deliberately designed the masking module to produce augmented graphs that posed challenge for GAT encoder to discriminate positive samples from negative samples, so that the graphs derived from same molecule were given similar embeddings but dissimilar embeddings from other molecules. Consequently, the contrastive learning model was forced to capture important chemical structure and higher-order semantic information. We validated the molecule representations obtained by contrastive learning on a couple of molecular property prediction tasks.

**Fig. 1.**
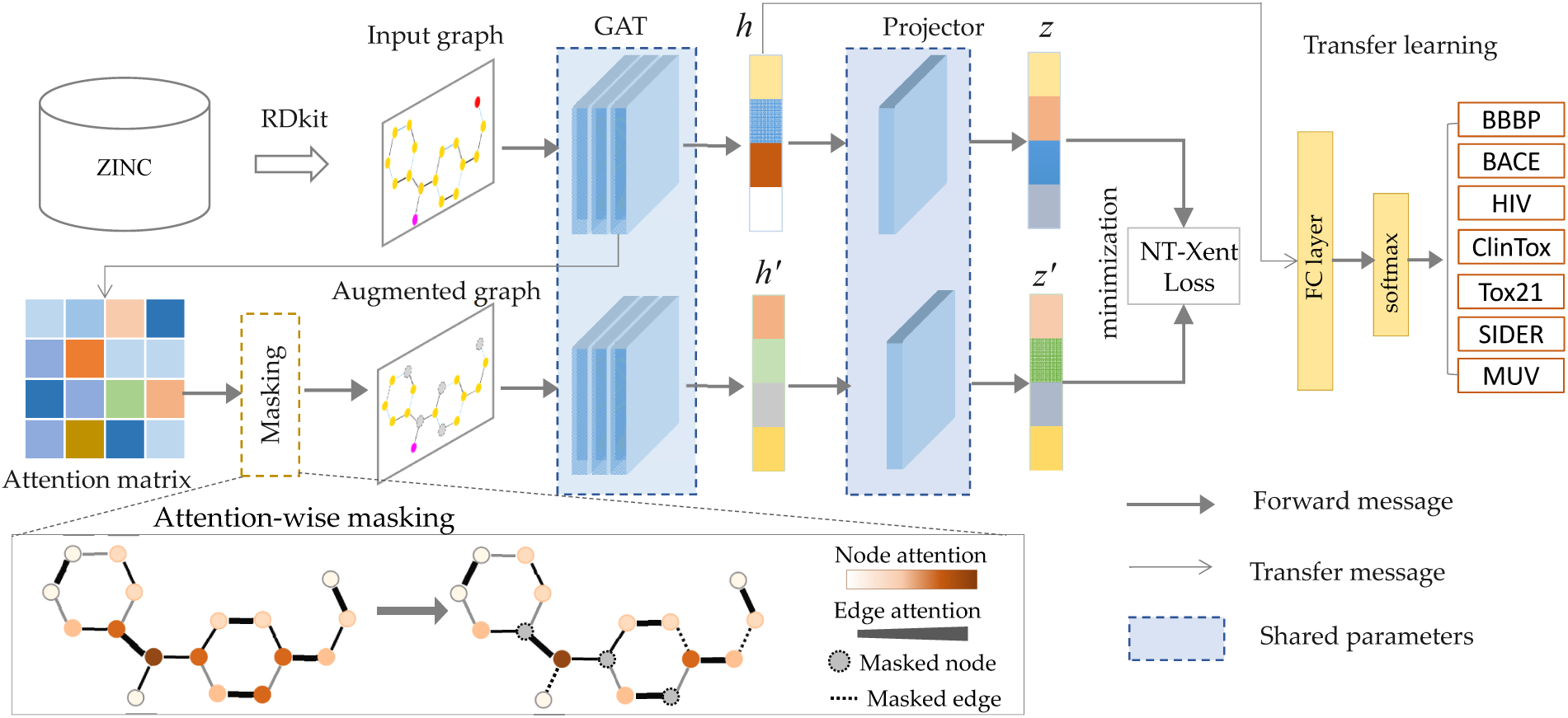
Illustrative flowchart of the proposed ATMOL contrastive learning framework for molecular property prediction.

### Molecule graph embedding

The graph attention network (GAT) is a multihead attention-based architecture developed to learn graph embedding. The GAT architecture was built from multiple graph attention layers, and each layer applied linear transformation to node-level representations for calculation of attention scores. Let *h*_*i*_ be the embedding of node *i, W* is a learnable attention weight matrix. The attention score *α*_*i,j*_ between node *i* and its first-order neighbor node *j* is calculated as

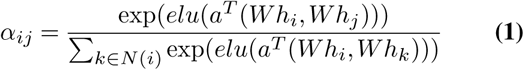

where *a* is a learnable vector, *elu* is the exponential linear unit activation function. The attention score *α*_*ij*_ was actually the *softmax* normalized message between node *i* and its neighbors. Once the attention scores were computed, the output feature of node *i* computed by aggregating its neighbor features weighted by corresponding attention scores:

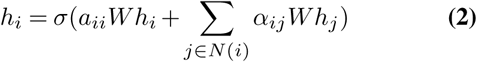

where *σ*(.) is the ReLU activation function. In our model, we used two GAT layers. The multi-head attention mechanism was applied to the first layer and the number of heads was set to 10. Given the node-level features, the global *max* pooling layer is used to obtain the graph embedding. The dimension of hidden features is set to 128. The multihead attention weight matrices in the first graph layer were aggregated into an integrative attention matrix in the second layer, which was used in the attention-wise mask module to produced augmentation graph.

### Attention-wise mask for graph augmentation

To produce high-quality augmented graph, we masked a percentage of nodes (edges) of the input molecule graph according to the attention scores learned by GAT encoder. When a node (edge) was masked, its embedding was set to 0 in the graph convolutional operation. If an edge was masked, the message passing along this edge was blocked. Formally, we define the masking rate *r*, representing the percent of nodes (edges) of the input graph would be masked. We iteratively masked nodes (edges) until the percentage of masked nodes (or edges) reached the predefined masking rate *r*. To explore the effect of masking different edges and nodes, we tried different masking strategies:

1. Max-attention masking: *r*% nodes (edges) with the largest attention scores were masked. This masking produced an augmented view with greatest difference to the current view of the input molecule graph.
2. Min-attention masking: *r*% nodes (edges) with the smallest attention scores were masked. This masking produced an augmented graph with least difference to the current view of the input molecule graph.
3. Random masking: *r*% nodes (edges) was randomly selected from the input graph and masked, neglecting the learned attention scores. This masking strategy was commonly used in previous studies, we thereby included it for comparison.
4. Roulette masking: the probability of each node or edge to be masked was proportional to its attention score. The attention weight matrix *W* was normalized by softmax function to yield a probability distribution. The possibility of an node (or edge) being masked is proportional to its probability.

### Contrastive learning

The GAT encoder transformed the input molecule graph and its augmented graph into embedding *h*_*i*_ and 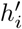, which were then mapped to *z*_*i*_ and 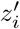 by a nonlinear projector. Next, the similarity 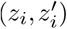 between two projected views was computed. We adopted the normalized temperature-scaled cross entropy (NT-Xent) as the contrastive loss function:

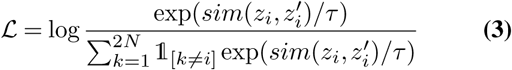

where 𝟙_[*k≠ i*]_ ∈ {0, 1} is an indicator function evaluating to 1 iff *k* ≠ *i, τ* denotes a temperature parameter; *N* is the number of samples in a mini-batch. In our study, the cosine distance was used to evaluate the similarity of two views came from one molecule.

The number and diversity of negative samples played crucial role in self-supervised representation learning, and previous studies have confirmed that a large number of negative samples could help improving performance. So, apart from the negative samples in a minibatch, we supplemented the augmented molecule graphs generated by attention-wise masking to the negative sample pool, so that the number of negative samples was greatly extended. Of more importance, the augmented molecule graph enriched the diversity of negative samples. The advantages our attention-wise masking for graph augmentation were reflected in two aspects. First, attention-wise masking generated challenging positive sample pairs, which increased the difficulty of contrastive learning and thereby prevented to learn collapsed latent representation. Also, it enriched the diversity of negative samples that were helpful for our model to learn molecular representations with good generalization.

### Pretraining and transfer learning

In the pretraining stage, the contrastive loss is minimized using the batch gradient descent algorithm by Adam optimizer. The learning rate is set to 1e-4, the batch-size is set to 128, and the number of pre-training epochs is set to 20 epochs.

During the transfer learning for molecular property prediction, we appended two fully-connected layers directly followed the GAT encoder. We froze the weights of GAT and tuned only the two fully-connected layers in the fine-tuning stage. The cross-entropy loss was applied for all classification task, and the learning rate is set to 1e-7. Adam optimizer was used and batch-size was set to 100. The performance of molecular property prediction was evaluated by ROC-AUC. The early stopping and dropout strategies were applied to prevent overfitting.

Each downstream dataset used for molecular property prediction was split into training, validation and test datasets in a ratio of 8:1:1. The transfer model was trained on the training set and validated on the validation set. To avoid random bias, the process was repeated for five times and each time evaluated on the test set. The mean AUC values were reported as the final performance.

## Results

### Contrastive learning boosted performance

We first verified whether contrastive learning-based pre-training improved the performance on downstream tasks. For this purpose, we compared our method to the model without pre-training on different molecular property prediction tasks. For each task, we trained a fully-supervised model, whose architecture consisted of a GAT encoder for molecular graph embedding and two fully-connected layers for molecular property prediction. To systematically evaluating the effectiveness of pre-training, we also took into account different molecular augmented graphs produced by masking nodes, edges or both nodes and edges (masking ratio r always was 25%). Table 2 showed the ROC-AUC values of these models on seven benchmark tasks. It can be seen that pre-training significantly boosted the performance on various molecular property prediction tasks.

**Table 2.**
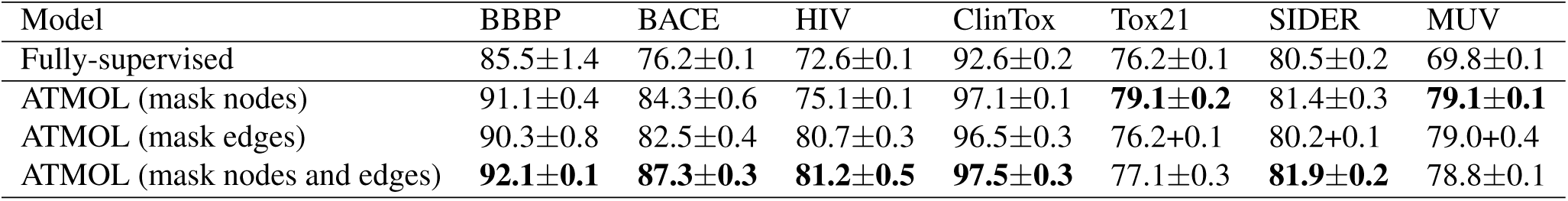
ROC-AUC (%) values of pretrained models and fully-supervised model on seven molecular property prediction tasks

| Model | BBBP | BACE | HIV | ClinTox | Tox21 | SIDER | MUV |
| --- | --- | --- | --- | --- | --- | --- | --- |
| Fully-supervised | 85.5±1.4 | 76.2±0.1 | 72.6±0.1 | 92.6±0.2 | 76.2±0.1 | 80.5±0.2 | 69.8±0.1 |
| ATMOL (mask nodes) | 91.1±0.4 | 84.3±0.6 | 75.1±0.1 | 97.1±0.1 | **79.1±0.2** | 81.4±0.3 | **79.1±0.1** |
| ATMOL (mask edges) | 90.3±0.8 | 82.5±0.4 | 80.7±0.3 | 96.5±0.3 | 76.2±0.1 | 80.2±0.1 | 79.0±0.4 |
| ATMOL (mask nodes and edges) | **92.1±0.1** | **87.3±0.3** | **81.2±0.5** | **97.5±0.3** | 77.1±0.3 | **81.9±0.2** | 78.8±0.1 |

Meanwhile, we found that the molecular graph augmented by masking both nodes and edges achieved best performance, compared to those augmented graphs by masking nodes or edges alone. The results showed that contrastive learning-based pre-training obtained informative molecular representations, and thereby greatly improved the performance of downstream tasks.

### Masking strategy affected feature extraction

To explore the influence of different graph augmentations on the feature extraction, we compared the performance derived from four masking strategies on seven downstream tasks. As shown in Figure 2, although the prediction performance varied on different molecular properties, the max-weight masking dominately achieved the best performance compared to other masking strategies. The random masking had the least performance. Seen by the GAT encoder, masking the nodes and edges with high attention scores produced an augmented graph that differed largely from the positive counterpart sample. So. we drew a conclusion that max-weight masking strategy posed a challenge for the contrastive learning to discriminate a pair of positive samples from a pool of negative samples came from other molecules. This challenge encouraged the model to learn informative molecular representations.

**Fig. 2.**
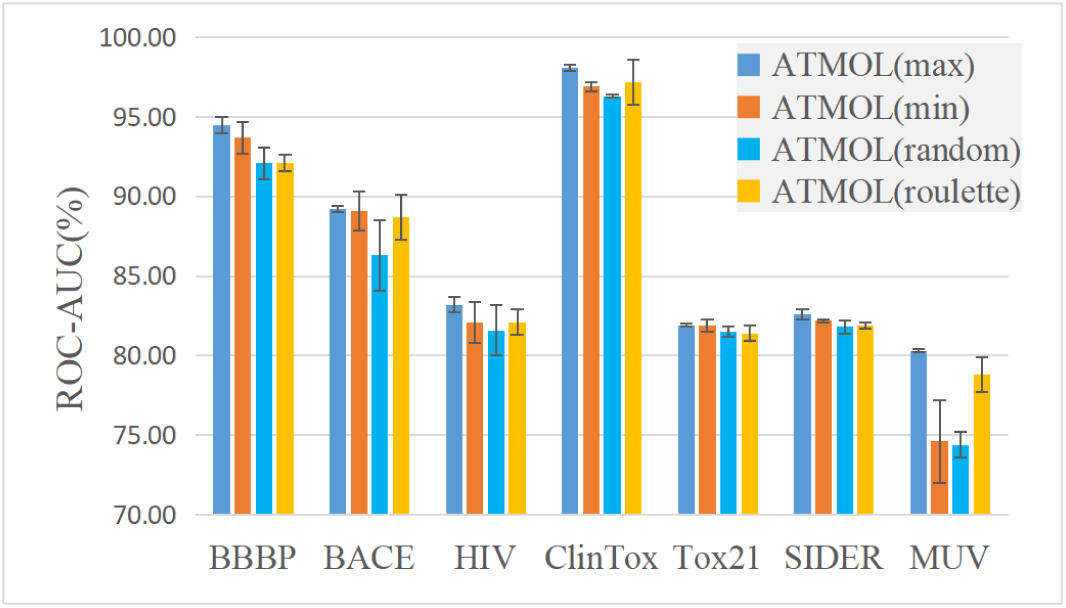
Performance comparison of four masking strategies for graph augmentation on seven molecular property prediction tasks

### Influence of masking rate

We went further to evaluate the impact of masking rate, namely, the percentage of masked nodes (edges). As max-weight masking has been shown to gain best performance, we evaluated its performance when different percentage edges were masked. As shown in Figure 3, the masking rate increased from 5% to 75%, the performance on seven downstream tasks rose up firstly, and reached the highest ROC-AUC values when masking rate was 25%. Thereafter, the performance decreased rapidly. Across all seven tasks, we observed similar tendency of performance. This result implied that too low masking rate did not produce effective augmented molecular graphs, while too high masking rate broke the essential chemical structure so that learned molecular representation was collapsed.

**Fig. 3.**
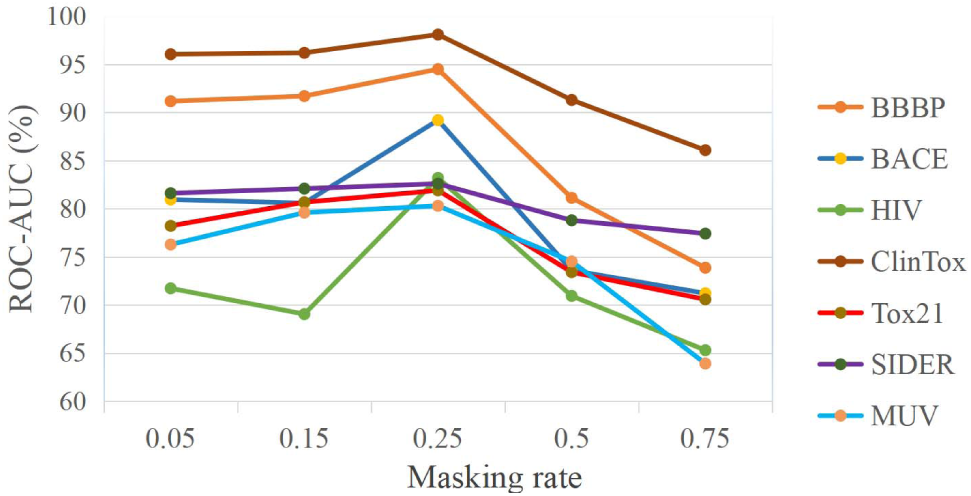
Performance achieved by different masking rate on seven downstream tasks

### Large-scale dataset improve representation learning

We were interested in whether larger scale of unlabeled data would improve the representation learning or not. So, apart from the *in vitro* set, we selected 3,000,000 molecules from the ZINC *now* set as another dataset, which was roughly tenfold of the *in vitro* set. For comparison, we referred them to as small and large set, respectively. We conducted representation learning on these two datasets separately, and then compared the performance on downstream tasks. As shown in Table 3, on the large dataset our model achieved higher performance over all downstream tasks. We concluded that self-supervised learning on large-scale dataset yielded molecular representations with better generalizablity.

**Table 3.** Performance comparison between small *vs*. large unlabeled datasets on seven molecular property prediction tasks

| Dataset | Masking strategy | BBBP | BACE | HIV | ClinTox | Tox21 | SIDER | MUV |
| --- | --- | --- | --- | --- | --- | --- | --- | --- |
| *in vitro*<br>(small) | Max-weight | **92.1±0.5** | **87.3±0.3** | **81.2±0.5** | **97.5±0.3** | **79.1±0.2** | **81.9±0.3** | **79.1±0.1** |
|  | Min-weight | 89.4±0.8 | 82.3±1.2 | 72.1±1.3 | 96.9±0.5 | 77.1±0.4 | 81.2±0.1 | 77.6±1.6 |
|  | random | 83.5±0.1 | 78.1±0.2 | 68.9±0.6 | 96.6±0.3 | 76.1±0.2 | 80.8±0.4 | 76.6±0.2 |
|  | roulette | 81.5±0.5 | 78.5±0.2 | 69.6±0.2 | 96.2±0.1 | 75.5±0.2 | 80.5±0.2 | 76.5±0.1 |
| *now subset*<br>(large) | Max-weight | **94.5±0.5** | **89.2±0.2** | **83.2±0.5** | **98.1±0.5** | **82.5±0.4** | **82.6±0.3** | **80.3±0.1** |
|  | Min-weight | 93.7±1.0 | 89.1±1.2 | 82.1±1.3 | 96.9±0.5 | 81.9±0.4 | 82.2±0.1 | 74.6±2.6 |
|  | random | 92.1±1.0 | 86.3±2.2 | 81.6±1.6 | 96.3±0.3 | 81.5±0.3 | 81.8±0.4 | 74.4±0.8 |
|  | roulette | 91.9±0.5 | 88.7±1.4 | 82.1±0.8 | 97.2±0.1 | 81.4±0.5 | 81.9±0.2 | 78.8±1.1 |

### Performance comparison to other methods

To verify the superior performance of our method, we compared it to five other competitive methods. All these methods used self-supervised learning for molecular feature extraction. We concisely introduced the methods as below:

- HU. et.al (33) pre-trained an expressive GNN at the level of individual nodes as well as entire graphs so as to learn useful local and global representations.
- N-Gram (34) run node embedding and then constructed a compact representation for the graph by assembling the node embeddings in short walks in the graph.
- GROVER (35) integrated GNN into Transformer with the context prediction task and the functional motif prediction task.
- MolCLR (27) proposed a graph contrast learning using graph neural network (GNNs), which generated contrastive pairs by randomly removal of nodes, edges or subgraphs.
- MGSSL (36) proposed topic-based graph self-supervised learning and a new GNN self-supervised topic generation framework.

Table 4 showed the ROC-AUC values of our method and five competitive methods on seven classification tasks. It was found that our method petrained on the *in vitro* dataset already outperformed five other methods on all tasks except MUV. Especially, on the SIDER prediction task, our method boosted the performance by 14% compared to the second best method MolCLR. Moreover, when pretrained on the large-scale dataset, our method achieved greater performance superiority. For example, on ClinTox and Tox21, our method outperformed all other methods by nearly 5%.

**Table 4.** ROC-AUC (%)of our method and five competitive methods on seven downstream tasks

| DataSet | BBBP | BACE | HIV | ClinTox | Tox21 | SIDER | MUV |
| --- | --- | --- | --- | --- | --- | --- | --- |
| HU. et.al[39] | 70.8±1.5 | 85.9±0.8 | 80.2±0.9 | 78.9±2.4 | 78.7±0.4 | 65.2±0.9 | 81.4±2.0 |
| N-Gram[40] | 91.2±3.0 | 87.6±3.5 | 83.0±1.3 | 85.5±3.7 | 76.9±2.7 | 63.2±0.5 | 81.6±1.9 |
| Grover[41] | 68.0±1.5 | 79.5±1.1 | 77.8±1.4 | 76.9±1.9 | 76.3±0.6 | 60.7±0.5 | 75.8±1.7 |
| MolCLR[42] | 73.6±0.5 | 89.0±0.3 | 80.6±1.1 | 93.2±1.7 | 79.8±0.7 | 68.0±1.1 | **88.6±2.2** |
| MGSSL[43] | 70.5±1.1 | 79.7±0.8 | 79.5±1.1 | 80.7±2.1 | 76.5±0.3 | 61.8±0.8 | 78.7±1.5 |
| ATMOL(small) | 92.1±0.5 | 87.3±0.3 | 81.2±0.5 | 97.5±0.3 | 79.1±0.2 | 81.9±0.3 | 79.1±0.1 |
| ATMOL(large) | **94.5±0.5** | **89.2±0.2** | **83.2±0.5** | **98.1±0.5** | **82.5±0.4** | **82.6±0.3** | 80.3±0.1 |

## Exploration of model interpretability

### Spatial localization of molecular representations

Spatial distribution of molecular representations was helpful to verify the effectiveness of the proposed method. We visualized the molecular representations before and after pretraining using UMAP tool (37), which is a manifold learning algorithm for dimension reduction with good preservation of data global structure. Figure 4 showed the 2D embeddings of molecules in BBBP and SIDER sets. The initial molecular representations spatially distributed in confusion, while after pretraining these molecules belonging to one same class gathered together and separated from other classes clearly. The observation illustrated the our method can effectively detect the physicochemical properties from chemical structure, so that the molecules with similar physicochemical properties gained similar latent representations.

**Fig. 4.**
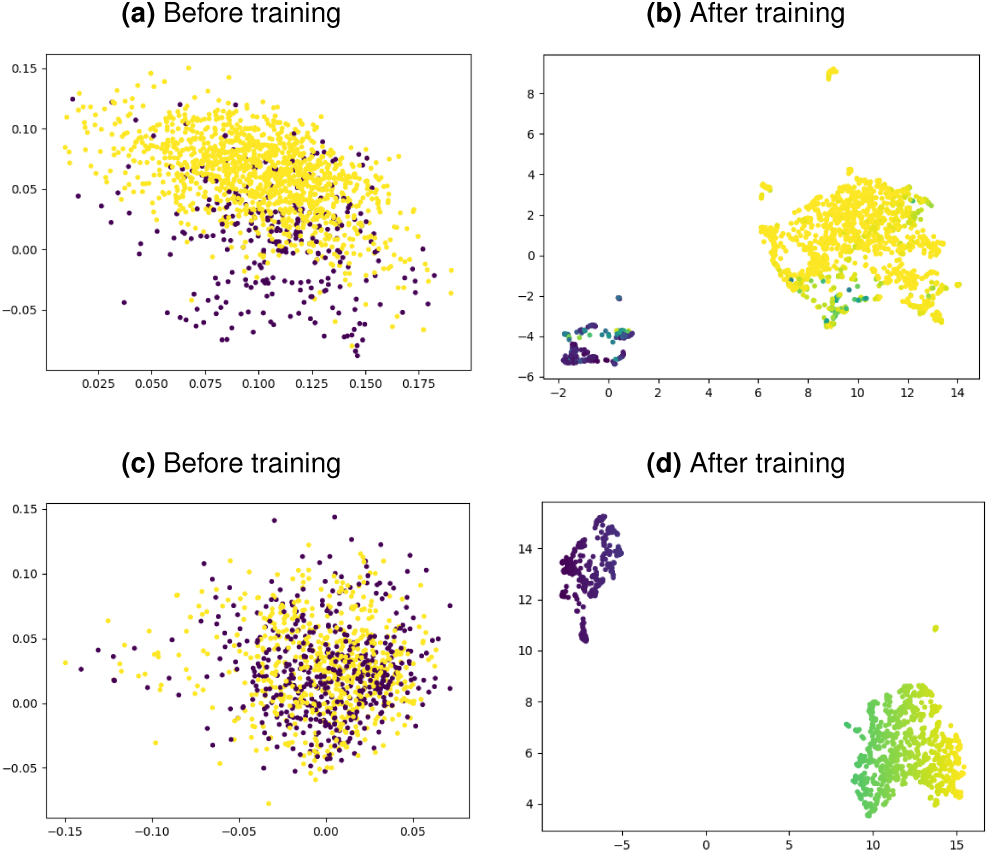
Visualization of spatial localization of molecular representations in BBBP and SIDER dataset. The left column showed the initial embeddings and the right column showed petrained embeddings.

### Attention weight revealed important chemical sub-structure

In the molecular representation learning stage, we tried to use graph attention network to identify chemical components that were important to specific prediction tasks. To investigate how the attention mechanism affected the focus of representation learning, we unfreeze the attention parameters and fine-tuned them during transfer learning. For intuitive interpretation, we visualized the attention weights in the molecular graph. From the BBBP dataset (38) regarding to membrane permeability, we randomly selected a molecule as an exemplar, whose SMILES is C[S](=O)(=O)c1ccc(cc1)[C@@H](O)[C@@H](CO) NC(=O)C(Cl)Cl. We computed the Pearson correlation coefficients of attention weights for each pair of atoms, and visualized the correlation matrix as a heatmap. As shown in Figure 5a, the heatmap displayed several highly correlated atomic groups, indicating that they functioned together to affect specific molecular properties. A further inspect can find that the benzene ring in this molecule may played important roles in determining the membrane permeability. Similarly, we randomly select a molecule FC1(F)COC(=NC1(C)c1cc(NC(=O)c2nn(cc2)C)ccc1F)N from the BACE dataset (39), its heatmap also illustrated a few significant atomic groups, as shown in Figure 5b.

**Fig. 5.**
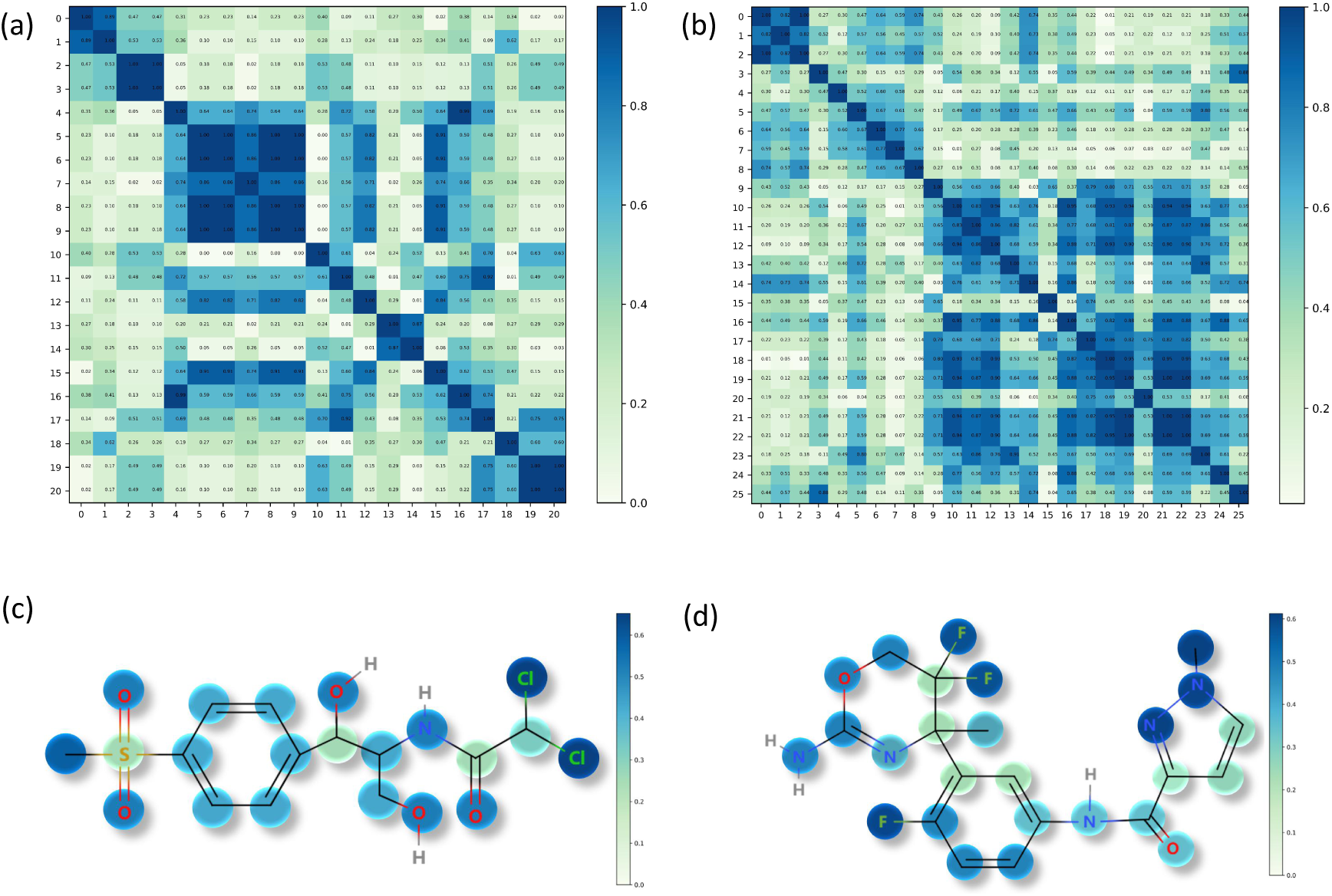
Visualization of correlation heatmaps of attention weights and molecule structure with atoms colored by attention weights. (a) and (c) showed the exemplar molecule selected from BBBP set, (b) and (d) showed the exemplar molecule selected from BACE set.

To further explore the influence of individual atoms on specific molecular properties, we visualized atomic-level attention weights. Keeping in mind the BBBP task focused on the membrane permeability, we found that two Cl atoms in the exemplar molecule had high attention weight, as shown in Figure 5c. Because the Cl atom has a strong electron attraction, we assume it affects the polarity of the molecule to a large extent, thereby affecting the membrane permeability. Also, since the hydroxyl groups promotes hydrophilicity, and we accordingly found that the hydroxyl groups were given relatively high attention weights (40).

Similarly, another exemplar molecule selected from BACE dataset is a human beta-secretase 1 (BACE-1) inhibitor. According to precious study by Mureddu et al (41), the heterocytosine aromatic family had the inhibitory effect to BACE-1. As shown in Figure 5d, the isocytosine component of this molecule has received more attention accordingly. The visualization and interpretability exploration illustrate how our model paid attention to relevant features from the perspective of molecular property prediction tasks.

## Discussion and Conclusion

The diversity of negative samples has been shown to greatly affect representation learning. There are two main methods to construct negative samples for contrastive learning. Some methods maintained a negative sample queue (22) and iteratively updated it in a FIFO manner, while other methods used only the samples in the current mini-batch as negative samples (23). In our study, beyond the samples in minibatch as negative samples, we added the augmented molecule graphs generated by attention-wise masking strategy to the negative sample queue, so that the negative samples was greatly extended and diversified.

Our attention-wise mask of molecular graph generated different contrastive views. By comparison, we found that the contrastive views generated by simultaneously masking edges and nodes achieved best performance in almost all down-stream tasks, except for Tox21 and MUV. This was consistent with the conclusions of MolCLR[28].

Moreover, we found that max-weight masking, i.e. masking the edges or nodes with large attention weights, achieved best performance. Intuitively, max-weight masking is similar to the idea of adversarial learning, by which each time an augmented molecular graph was generated with the largest difference to the positive couterpart view, from the perspective of contrastive loss. Moreover, the graph augmentation process dynamically tracked the change of attention weights, so that our model was forced to inspect different components of the molecular graph and finally reach steady state. Therefore, we concluded that the challenging contrastive views were helpful to learn important semantic structure.

In summary, our self-supervised representation learning on large-scale unlabeled molecules significantly improved the performance of various molecular property prediction tasks. This a task-agnostic pretraining and thus yielded the molecular representation with desirable expressiveness and generalizability.

## ACKNOWLEDGEMENTS

This work was supported by the National Natural Science Foundation of China under grants No. 62072058 and No. 61972422.

## Bibliography

1. M. Axelrod, V. L. Gordon, M. Conaway, A. Tarcsafalvi, D. J. Neitzke, D. Gioeli, and M. J. Weber. Combinatorial drug screening identifies compensatory pathway interactions and adaptive resistance mechanisms. Oncotarget, 4(4):622–35, 2013.

2. C. M. Song, S. J. Lim, and J. C. Tong. Recent advances in computer-aided drug design. Brief Bioinform, 10(5):579–91, 2009.

3. A. P. Bartok, R. Kondor, and G. Csanyi. On representing chemical environments. Physical Review B, 87(18), 2013. ISSN 2469-9950.

4. L. M. Ghiringhelli, J. Vybiral, S. V. Levchenko, C. Draxl, and M. Scheffler. Big data of materials science: critical role of the descriptor. Phys Rev Lett, 114(10):105503, 2015.

5. L. David, A. Thakkar, R. Mercado, and O. Engkvist. Molecular representations in ai-driven drug discovery: a review and practical guide. Journal of Cheminformatics, 12(1), 2020. ISSN 1758-2946.

6. R. Bade, H. F. Chan, and J. Reynisson. Characteristics of known drug space. natural products, their derivatives and synthetic drugs. European Journal of Medicinal Chemistry, 45 (12):5646–5652, 2010. ISSN 0223-5234.

7. A. Cereto-Massague, M. J. Ojeda, C. Valls, M. Mulero, S. Garcia-Vallve, and G. Pujadas. Molecular fingerprint similarity search in virtual screening. Methods, 71:58–63, 2015.

8. D. Rogers and M. Hahn. Extended-connectivity fingerprints. J Chem Inf Model, 50(5):742–54, 2010.

9. S. J. Pan, I. W. Tsang, J. T. Kwok, and Q. A. Yang. Domain adaptation via transfer component analysis. Ieee Transactions on Neural Networks, 22(2):199–210, 2011.

10. Kaiming He, Haoqi Fan, Yuxin Wu, Saining Xie, and Ross Girshick. Momentum contrast for unsupervised visual representation learning. In Proceedings of the IEEE/CVF conference on computer vision and pattern recognition, pages 9729–9738, 2020.

11. Thomas N Kipf and Max Welling. Semi-supervised classification with graph convolutional networks. arXiv preprint arXiv:1609.02907, 2016.

12. D. Duvenaudt, D. Maclaurin, J. Aguilera-Iparraguirre, R. Gomez-Bombarelli, T. Hirzel, A. Aspuru-Guzik, and R. P. Adams. Convolutional networks on graphs for learning molecular fingerprints. Advances in Neural Information Processing Systems 28 (Nips 2015), 28, 2015. ISSN 1049-5258.

13. J. Gilmer, S. S. Schoenholz, P. F. Riley, O. Vinyals, and G. E. Dahl. Neural message passing for quantum chemistry. International Conference on Machine Learning, Vol 70, 70, 2017. ISSN 2640-3498.

14. M. Karamad, R. Magar, Y. T. Shi, S. Siahrostami, I. D. Gates, and A. B. Farimani. Orbital graph convolutional neural network for material property prediction. Physical Review Materials, 4(9), 2020. ISSN 2475-9953.

15. S. Chmiela, H. E. Sauceda, K. R. Muller, and A. Tkatchenko. Towards exact molecular dynamics simulations with machine-learned force fields. Nature Communications, 9, 2018. ISSN 2041-1723.

16. V. L. Deringer, N. Bernstein, A. P. Bartok, M. J. Cliffe, R. N. Kerber, L. E. Marbella, C. P. Grey, S. R. Elliott, and G. Csanyi. Realistic atomistic structure of amorphous silicon from machine-learning-driven molecular dynamics. Journal of Physical Chemistry Letters, 9(11):2879–2885, 2018. ISSN 1948-7185.

17. W. J. Wang and R. Gomez-Bombarelli. Coarse-graining auto-encoders for molecular dynamics. Npj Computational Materials, 5(1), 2019.

18. H. Altae-Tran, B. Ramsundar, A. S. Pappu, and V. Pande. Low data drug discovery with one-shot learning. ACS Cent Sci, 3(4):283–293, 2017.

19. H. Chen, O. Engkvist, Y. Wang, M. Olivecrona, and T. Blaschke. The rise of deep learning in drug discovery. Drug Discov Today, 23(6):1241–1250, 2018.

20. Z. K. Hao, C. Q. Lu, Z. Y. Huang, H. Wang, Z. Y. Hu, Q. Liu, E. H. Chen, and C. Lee. Asgn: An active semi-supervised graph neural network for molecular property prediction. Kdd ‘20: Proceedings of the 26th Acm Sigkdd International Conference on Knowledge Discovery & Data Mining, pages 731–739, 2020.

21. J. Yosinski, J. Clune, Y. Bengio, and H. Lipson. How transferable are features in deep neural networks ? Advances in Neural Information Processing Systems 27 (Nips 2014), 27, 2014.

22. Ting Chen, Simon Kornblith, Kevin Swersky, Mohammad Norouzi, and Geoffrey E Hinton. Big self-supervised models are strong semi-supervised learners. Advances in neural information processing systems, 33:22243–22255, 2020.

23. T. Chen, S. Kornblith, M. Norouzi, and G. Hinton. A simple framework for contrastive learning of visual representations. International Conference on Machine Learning, Vol 119, 119, 2020. ISSN 2640-3498.

24. Jacob Devlin, Ming-Wei Chang, Kenton Lee, and Kristina Toutanova. Bert: Pre-training of deep bidirectional transformers for language understanding, October 01, 2018 2018.

25. X. C. Zhang, C. K. Wu, Z. J. Yang, Z. X. Wu, J. C. Yi, C. Y. Hsieh, T. J. Hou, and D. S. Cao. Mg-bert: leveraging unsupervised atomic representation learning for molecular property prediction. Brief Bioinform, 22(6), 2021.

26. Viraj Bagal, Rishal Aggarwal, PK Vinod, and U Deva Priyakumar. Molgpt: Molecular generation using a transformer-decoder model. Journal of Chemical Information and Modeling, 2021. ISSN 1549-9596.

27. Yuyang Wang, Jianren Wang, Zhonglin Cao, and Amir Barati Farimani. Molclr: molecular contrastive learning of representations via graph neural networks. arXiv preprint arXiv:2102.10056, 2021.

28. Yifan Hou, Jian Zhang, James Cheng, Kaili Ma, Richard TB Ma, Hongzhi Chen, and Ming-Chang Yang. Measuring and improving the use of graph information in graph neural networks. In International Conference on Learning Representations, 2020.

29. P. Li, J. Wang, Y. Qiao, H. Chen, Y. Yu, X. Yao, P. Gao, G. Xie, and S. Song. An effective self-supervised framework for learning expressive molecular global representations to drug discovery. Brief Bioinform, 22(6), 2021.

30. Greg Landrum. Rdkit: A software suite for cheminformatics, computational chemistry, and predictive modeling, 2013.

31. Z. Wu, B. Ramsundar, E. N. Feinberg, J. Gomes, C. Geniesse, A. S. Pappu, K. Leswing, and V. Pande. Moleculenet: a benchmark for molecular machine learning. Chem Sci, 9(2): 513–530, 2018.

32. Bharath Ramsundar, Peter Eastman, Patrick Walters, and Vijay Pande. Deep learning for the life sciences: applying deep learning to genomics, microscopy, drug discovery, and more. O’Reilly Media, 2019. ISBN 1492039802.

33. Weihua Hu, Bowen Liu, Joseph Gomes, Marinka Zitnik, Percy Liang, Vijay Pande, and Jure Leskovec. Strategies for pre-training graph neural networks. arXiv preprint arXiv:1905.12265, 2019.

34. Shengchao Liu, Mehmet F Demirel, and Yingyu Liang. N-gram graph: Simple unsupervised representation for graphs, with applications to molecules. Advances in neural information processing systems, 32, 2019.

35. Yu Rong, Yatao Bian, Tingyang Xu, Weiyang Xie, Ying Wei, Wenbing Huang, and Junzhou Huang. Self-supervised graph transformer on large-scale molecular data. Advances in Neural Information Processing Systems, 33:12559–12571, 2020.

36. Zaixi Zhang, Qi Liu, Hao Wang, Chengqiang Lu, and Chee-Kong Lee. Motif-based graph self-supervised learning for molecular property prediction. Advances in Neural Information Processing Systems, 34, 2021.

37. Leland McInnes, John Healy, and James Melville. Umap: uniform manifold approximation and projection for dimension reduction. 2020.

38. I. F. Martins, A. L. Teixeira, L. Pinheiro, and A. O. Falcao. A bayesian approach to in silico blood-brain barrier penetration modeling. J Chem Inf Model, 52(6):1686–97, 2012.

39. Govindan Subramanian, Bharath Ramsundar, Vijay Pande, and Rajiah Aldrin Denny. Computational modeling of β-secretase 1 (bace-1) inhibitors using ligand based approaches. Journal of chemical information and modeling, 56(10):1936–1949, 2016.

40. Yin Fang, Qiang Zhang, Haihong Yang, Xiang Zhuang, Shumin Deng, Wen Zhang, Ming Qin, Zhuo Chen, Xiaohui Fan, and Huajun Chen. Molecular contrastive learning with chemical element knowledge graph. arXiv preprint arXiv:2112.00544, 2021.

41. Luca G Mureddu and Geerten W Vuister. Fragment-based drug discovery by nmr. where are the successes and where can it be improved. Frontiers in molecular biosciences, page 110, 2022. ISSN 2296-889X.

